# Distributed Code for Semantic Relations Predicts Neural Similarity

**DOI:** 10.1101/596726

**Authors:** Jeffrey N. Chiang, Yujia Peng, Hongjing Lu, Keith J. Holyoak, Martin M. Monti

## Abstract

The ability to generate and process semantic relations is central to many aspects of human cognition. Theorists have long debated whether such relations are coded as atomistic links in a semantic network, or as distributed patterns over some core set of abstract relations. The form and content of the conceptual and neural representations of semantic relations remains to be empirically established. The present study combined computational modeling and neuroimaging to investigate the representation and comparison of abstract semantic relations in the brain. By using sequential presentation of verbal analogies, we decoupled the neural activity associated with encoding the representation of the first-order semantic relation between words in a pair from that associated with the second-order comparison of two relations. We tested alternative computational models of relational similarity in order to distinguish between rival accounts of how semantic relations are coded and compared in the brain. Analyses of neural similarity patterns supported the hypothesis that semantic relations are coded, in the parietal cortex, as distributed representations over a pool of abstract relations specified in a theory-based taxonomy. These representations, in turn, provide the immediate inputs to the process of analogical comparison, which draws on a broad frontoparietal network. This study sheds light not only on the form of relation representations but also on their specific content.

**Significance:** Relations provide basic building blocks for language and thought. For the past half century, cognitive scientists exploring human semantic memory have sought to identify the code for relations. In a neuroimaging paradigm, we tested alternative computational models of relation processing that predict patterns of neural similarity during distinct phases of analogical reasoning. The findings allowed us to draw inferences not only about the form of relation representations, but also about their specific content. The core of these distributed representations is based on a relatively small number of abstract relation types specified in a theory-based taxonomy. This study helps to resolve a longstanding debate concerning the nature of the conceptual and neural code for semantic relations in the mind and brain.

## Introduction

The poet Samuel Taylor Coleridge claimed that the creative mind needs to become “accustomed to contemplate not *things* only, … but likewise and chiefly the *relations* of things….” (*1*) Because relations provide basic building blocks for language and thought, they are central for a range of cognitive tasks. A prime example is the critical role of relation representations in analogical reasoning (*2*), a mental process that impacts human activities as diverse as metaphor comprehension (*3*), mathematics education (*4*), scientific discovery (*5*) and engineering design (*6*). But while the importance of relations is widely recognized, no consensus has emerged regarding the form of relation representations in the mind and brain.

For the past half century, cognitive scientists exploring human semantic memory have sought to identify the nature of the code for relations (for reviews see Refs. (*7, 8*)). Two longstanding views, mainly based on data from speeded verification of category-membership relations (e.g., deciding as rapidly as possible whether a rose *is a* flower), continue to be influential. One approach, originating in computer science (*9*), treats relations as labeled unitary links between localist nodes representing concepts (e.g., an “*is a*” link connecting rose to flower). Relation verification is viewed as an all-or-none process of retrieving the relevant link. Current computational models of analogy based on traditional symbolic knowledge representations (*10*) continue to assume relations are coded as atomistic links. In contrast, an alternative view hypothesizes that rather than being represented by explicit links, relations between concepts are computed by operations performed on featural representations of concepts (*11, 12*). In support of the latter view, analyses of verification time based on speed-accuracy decomposition have revealed that relation information accrues continuously over time, rather than being retrieved in an all-or-none fashion (*13, 14*). These findings suggest that relations are not stored and retrieved as atomic units. However, the specific nature and content of relation representations remain a mystery. There is continuing debate as to whether a relation is coded explicitly (i.e., each relation has its own semantic representation), or whether a relation is coded only implicitly based on the features of the concepts it links (*15*). The resolution of this debate is likely to require evidence that goes beyond behavioral measures such as verification times.

Here we test a model of relation representation, combining recent advances in machine learning and cognitive science with neuroimaging. Our tests are based on a key property of distributed representations: they predict systematic and graded variations in similarity, rather than simply an all-or-none distinction between same and different. We propose that specific semantic relations between words are coded as distributed representations over a set of abstract relations, specified in a taxonomy founded on linguistic and psychological evidence (*16*). This taxonomy includes ten general types of relations (e.g., *similar, contrast, cause-purpose*), each of which has several subtypes, resulting in a total of 79 semantic relations. After a computational model has acquired knowledge of these relations, the specific relation between any pair of words can be represented as a vector based on the posterior probability that the pair instantiates each known relation. The resulting distributed representation captures the intuition that many word pairs instantiate multiple relations to some degree. For example, the concepts *hill-mountain* primarily instantiate the relation of *similar* (both are types of high geological formations), but they also to some degree instantiate *contrast* (differing in height).

Following Coleridge (*1*), to represent relations between things it is first necessary to have representations of those “things”—in the case of semantic relations, we first need semantic representations of individual words. To represent word meanings, we adopt word embeddings produced by a recent machine-learning model, Word2vec (*17*). This model applies a predictive learning algorithm to a large text corpus (e.g., Google News) to create high-dimensional semantic vectors for individual words. Vectors generated by Word2vec and similar models have been show to accurately capture human judgments of semantic similarities among words (*18*), and have also been used to create a neural decoder to predict patterns of brain activity produced in response to sentences (*19*).

Using semantic vectors for individual words derived by Word2vec, pairs of words can be used to learn representations of individual relations in the taxonomy using a model called *Bayesian Analogy with Relational Transformations* (*BART*; see Refs. (*20, 21*)). BART is trained with a small number of word pairs (∼20 pairs) as positive examples of each specific relation in the taxonomy (*16*). After learning the set of 79 abstract relations, for any word pair BART can estimate the probability that the word pair instantiates each learned relation. The model accurately predicts human judgments of how well word pairs instantiate relations (i.e., relation prototypicality)(*21*).

When the entire set of learned relations is considered, any word pair can be coded as a vector of relation probabilities, which constitutes a distributed representation of the specific relation between the two words (Figure 1). The high-dimensional input vectors for individual words are thus transformed into a new (and much more interpretable) vector space that codes the semantic relations between words. BART’s relation vectors enable computations of second-order relational similarity between word pairs, providing a direct basis for solving verbal analogies in the form *A:B*::*C:D* (e.g., *old:young* :: *hot:cold*). To evaluate an analogy between two word pairs, BART assesses their second-order relation similarity based on the cosine distance between the two distributed patterns of relations (with low cosine values indicative of a good analogical match). Behavioral evidence indicates that BART can solve a set of simple verbal analogies with a degree of accuracy comparable to humans (*21*).

**Figure 1.**
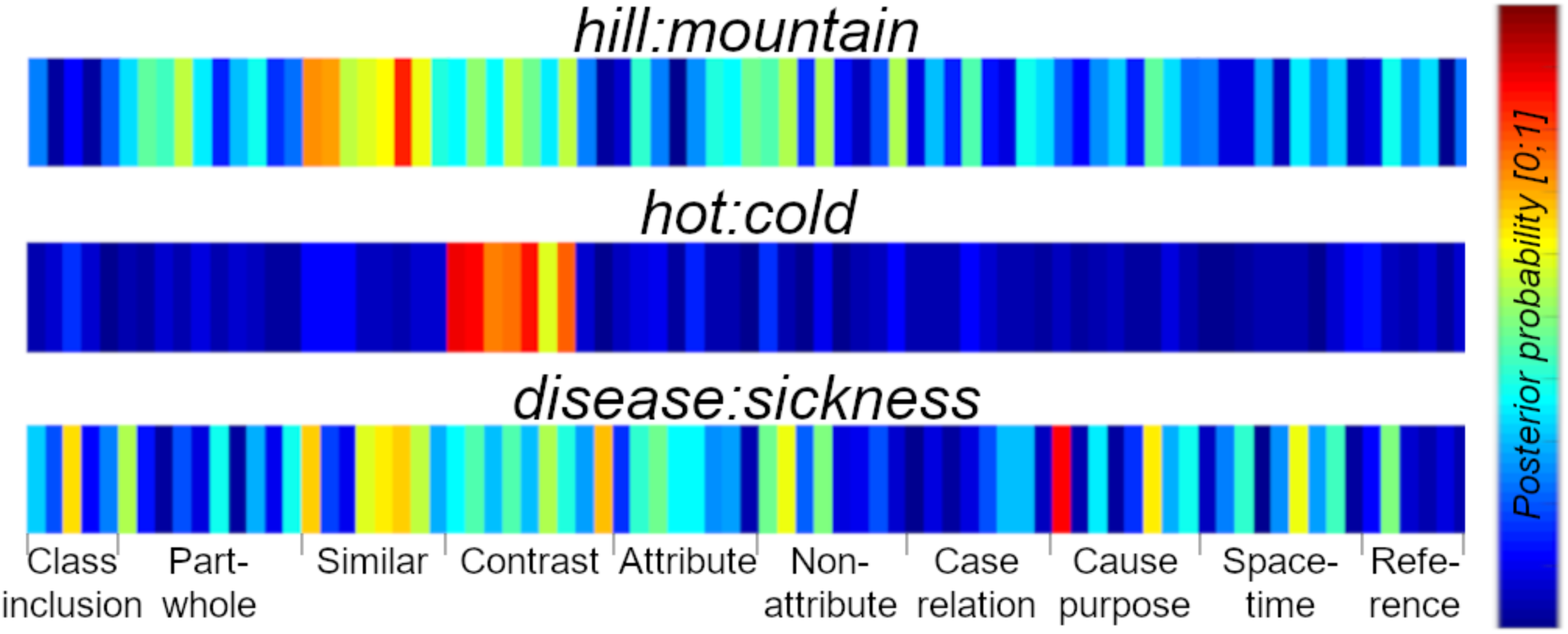
Examples of distributed representations of the relation instantiated by an individual word pair, generated by the BART model (*21*). Each word pair primarily instantiates one of the three general relation types used in the neuroimaging experiment reported here: *hill-mountain* (*similar*), *hot-cold* (*contrast*), *disease-sickness* (*cause-purpose*). The distributed code is based on the posterior probabilities that a given pair instantiates each of 79 abstract relations, drawn from 10 major types (*16*). The representations vary in their degree of distribution across the pool of abstract relations, with each word pair attaining its highest value on one or more relations within its general type.

To test the proposed distributed code for semantic relations, we performed a study of verbal analogical reasoning using functional magnetic resonance imaging (fMRI) to measure similarities among neural responses during different stages of relational processing. The basic logic was to use BART and baseline models to predict degrees of relation similarity, and then to correlate predictions of the models with patterns of neural similarity in regions of interest. We examined three types of abstract relations (*similar, contrast, cause-purpose*), with three specific relations for each type (see *Methods and Materials*). For each relation, we selected 16 word pairs high in typicality as assessed by human judgments (*22*), yielding 48 word pairs per relation type. Using the resulting 144 (48 examples × 3 relation types) distinct word pairs, we formed pairs of pairs to create verbal analogy problems in the form *A:B* :: *C:D* (valid) or else *A:B* ::*C’:D’* (invalid).

We employed a sequential event-related fMRI design (23) to separate the construction of first-order relations (i.e., relations between words in a pair) from the second-order assessment of similarity between relations (i.e., the analogical match between *A:B* and *C:D* relations) (see Figure 2; also *Materials and Methods*). The *A:B* phase provides a relatively pure measure of neural activity involved in coding the individual *A* and *B* words and the *A:B* relation. The *C:D* phase includes the neural computation required to *compare* the two relations (as well as neural activity required to maintain the *A:B* relation and to represent the *C:D* relation). If semantic relations have distributed representations based on the taxonomy of abstract relations, we should find brain regions in which BART is the best predictor of neural similarity. In contrast, if relations are coded as atomic units, then similarity of two word pairs will only depend on whether they instantiate the same or different relation types.

**Figure 2.**
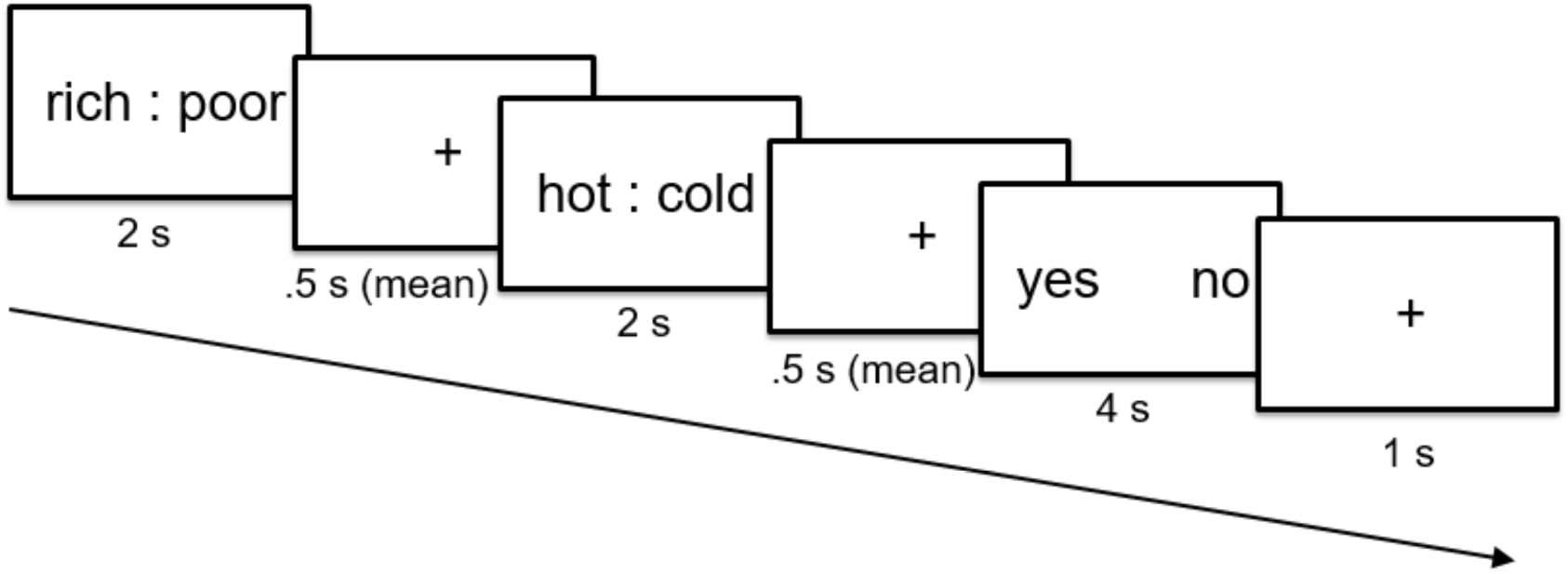
Timing of events on each trial. Participants were shown two word pairs, first an *A:B* pair for 2 seconds, then a *C:D* pair for 2 seconds after a jitter, and finally a cue to make a yes/no decision about the validity of the analogy. Participants responded by pressing a button box, where the location of “yes” and “no” buttons varied from trial to trial, making it impossible to plan a specific motor response until the first two phases had been completed. In a rapid event-related fMRI design, healthy young adults were asked to evaluate two pairs of semantic concepts. Each analogy was presented as two pairs of words, an *A:B* pair (e.g., *rich:poor*) followed by a *C:D* pair (e.g., *hot:cold*) exemplifying the same relation (here *contrast*) as the *A:B* pair (valid analogy), or else a *C’:D’* pair (e.g., *loss:grief*) exemplifying a different relation (here *cause-purpose*) (invalid analogy). All analogies were based on word pairs taken from a set of norms (22), exemplifying three abstract relation types (*similar, contrast*, and *cause-purpose*) (16) [see *supporting information (SI) Appendix*, Table S4].

**Figure 3.**
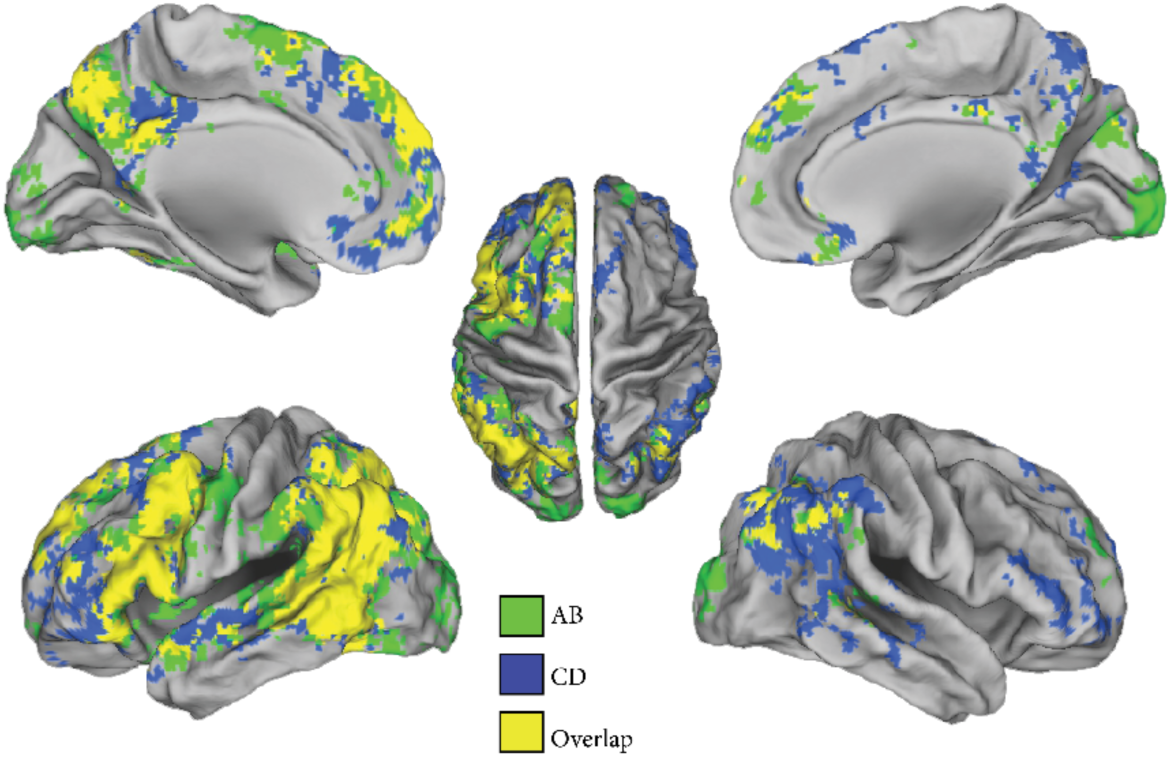
MVPA searchlight results. Regions in which the three general semantic relations could be discriminated above chance during different phases of the analogy task (*p* < 0.01, corrected for multiple comparisons using TFCE cluster correction (*28*)).

## Results

In what follows we describe a three-pronged approach to characterizing the computational architecture supporting first- and second-order relational processing. First, we use a multivariate pattern analysis (MVPA; (*24*)) approach to confirm the sensitivity of large, known, regions of cortex to different types of semantic relations. Second, we employ a model-guided representational similarity analysis (RSA;(*25, 26*)) approach to test the neural plausibility of the first-order (i.e., *A:B*) relational representations described by four possible computational accounts of semantic relations. Finally, we employ a second model-guided analysis to test at the neural level the predicted patterns of second-order (i.e., *A:B::C:D*) relational comparisons suggested by different computational accounts.

### Decoding Neural Activity Patterns to Classify Relation Types

To characterize the representations of abstract semantic relations in the brain, we first conducted a multivariate pattern analysis using a searchlight method (*27*). This analysis revealed left-lateralized areas of the brain capable of distinguishing different types of abstract relations on the basis of activation patterns across both the *A:B* and *C:D* phases. In addition, during the second-order comparison (i.e., the *C:D* phase) the three abstract relations could also be distinguished in left rostrolateral and right fronto-temporal cortices. (See *SI Appendix*, Supplemental Data Analyses, for a detailed report and also for a univariate analyses of areas active during the *A:B* phase versus rest, *C:D* phase versus rest, and *C:D* versus *A:B*.)

### Comparing Computational Accounts of First-order Relational Similarity (*A:B Phase*)

The initial decoding analysis established that the three relation types can be classified using brain activity patterns. However, that analysis treats each word pair as instantiating a single relation type, generating only a coarse measure of similarity (same or different relation type). To assess more detailed computational accounts that predict graded similarity patterns, we examined item-level similarity across word pairs using Representational Similarity Analysis (RSA; (*25, 26, 29*)).

Theoretical patterns of dissimilarity across word pairs were derived from four computational models. First, a design matrix based on the three relation types was used as a baseline model, which predicts relation similarity is all-or-none (i.e., same versus different relation types). Two additional control models were tested, both of which derive similarity predictions directly from Word2vec vectors for the individual words in a pair. These two control models differ in their assumptions about how (or whether) the relation between the two words is represented. Under *Word2vec-concat*, the meaning of the words within a pair is a simple aggregate of the semantic vectors of the two individual words. The similarity between any two word pairs is computed by the cosine distance between the two concatenated vectors. This model is nonrelational, instead capturing semantic similarity across pairs based solely on the meanings of the individual words. Word2vec-concat serves to identify patterns of similarity based on lexical semantics, separate from any representation of the relation between the two words within each pair. Under *Word2vec-diff*, the first-order relation between two words is defined in a generic fashion as the *difference* between the semantic vectors of each word within a pair; second-order similarity of relations is assessed by the cosine distance between the two difference vectors that form the analogy. This model, which has been directly applied to analogy problems in work on machine learning (*18*), codes relations only implicitly (i.e., as a difference vector computed from individual words).

The fourth model, BART, creates a distributed representation of the specific relation between a pair of words based on the posterior probabilities that the word pair instantiates each relation in a theory-based taxonomy (*16*). Unlike any of the control models, BART assumes that the specific relation between a pair of words has an explicit distributed representation. The similarity between any two word pairs is computed by the cosine distance between the two relation vectors. As illustrated in Figure 4, BART and the two control models based on Word2vec each take identical inputs (Word2vec vectors for the individual words in a pair) and assume an identical computation for relation similarity between two word pairs (cosine distance between the final vector produced by the model) (see *SI Appendix*, Supplemental Details for Computational Models).

**Figure 4.**
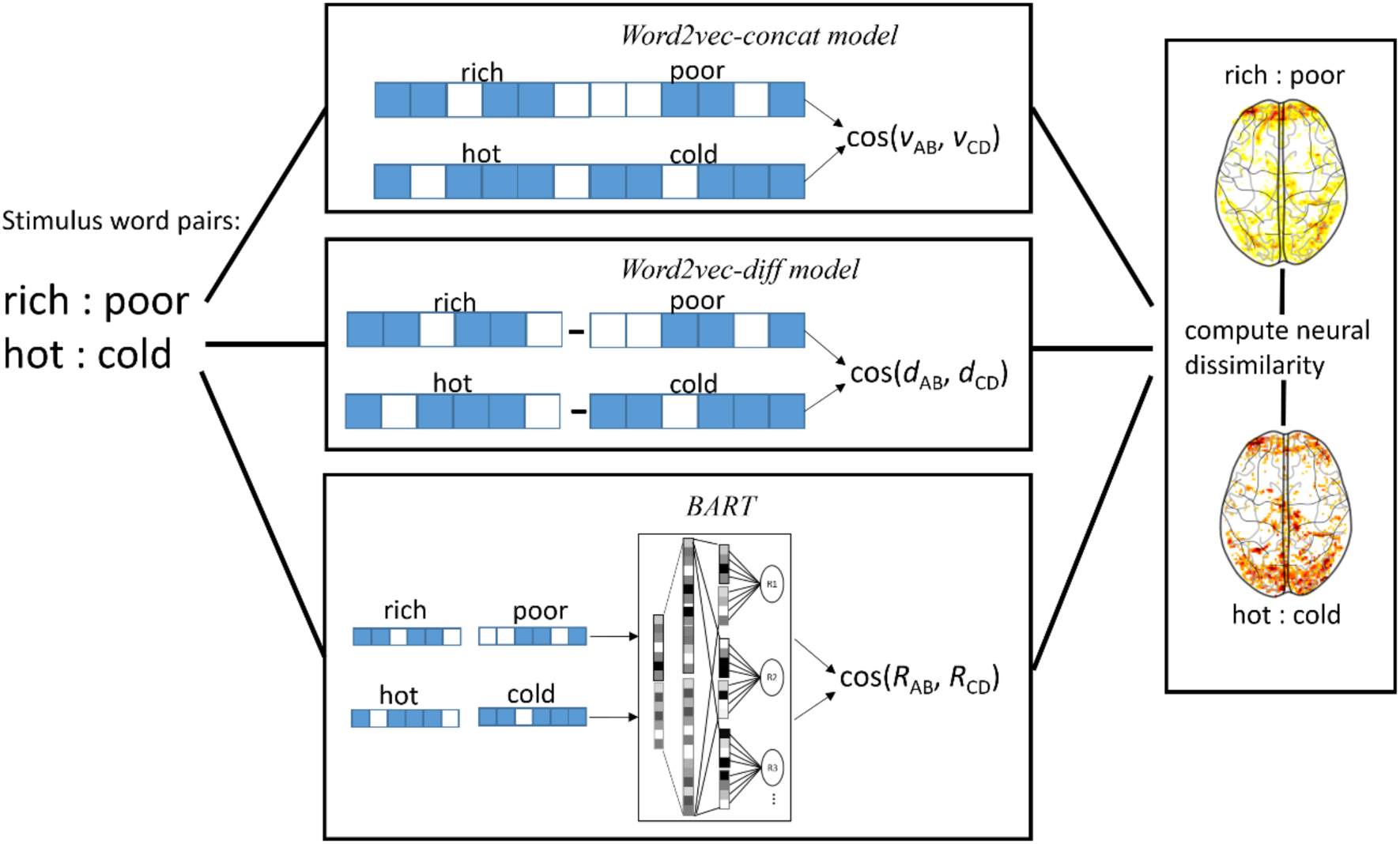
Model-guided approach to discovering neural signatures of specific relations. For any two word pairs (e.g., *rich:poor, hot:cold*), three alternative models are used to predict dissimilarity based on the cosine distance between the representations of each individual word pair, using 300-dimensional Word2vec vectors as inputs (left). *Word2vec-concat* (nonrelational) concatenates the vectors for individual words in a pair; *Word-2vec-diff* (generic relation) defines the relation as the difference vector; *BART* (specific relations) creates a new relational vector for each pair based on previously-learned relations. The neural response to each word pair (right) is obtained, allowing a calculation of dissimilarity between patterns of voxels. Neural dissimilarities are compared with computational predictions in order to arbitrate between alternative models.

Using a searchlight procedure, Representational Dissimilarity Matrices (RDMs) predicted by each of the four models (relation-types baseline, BART, Word2vec-concat, Word2vec-diff) were compared to the empirical pattern of dissimilarity across word pairs observed in neural activity patterns during the *A:B* phase (see *SI Appendix*, Representational Similarity Analysis). These analyses were conducted at the level of individual word pairs; hence the size of each RDM was 144 × 144 (Figure 5). Among the four models that were tested, only the RDM derived from BART yielded significant correlations with neural RDMs (Figure 6A). These correlations primarily involved the left superior parietal lobe (lSPL) and intraparietal sulcus (lIPS). In approximately the same regions, the correlation for BART was significantly greater than those for either of the other computational models (Figure 6B) or for the relation-types baseline model (Figure 6C). These findings indicate that the distributed representation of relations postulated by BART is realized in regions within the posterior cortex previously reported to be involved in the formation and/or retrieval of relational information (*30*–*32*). Notably, neither BART nor any of the other models yielded significant correlations with the neural patterns of activity in rostrolateral PFC (Brodmann area (BA)10m, BA10l), consistent with the hypothesis that the RLPFC is not involved in representing first-order relations, but rather is primarily engaged in making second-order *comparisons* between relations (*33, 34*).

**Figure 5.**
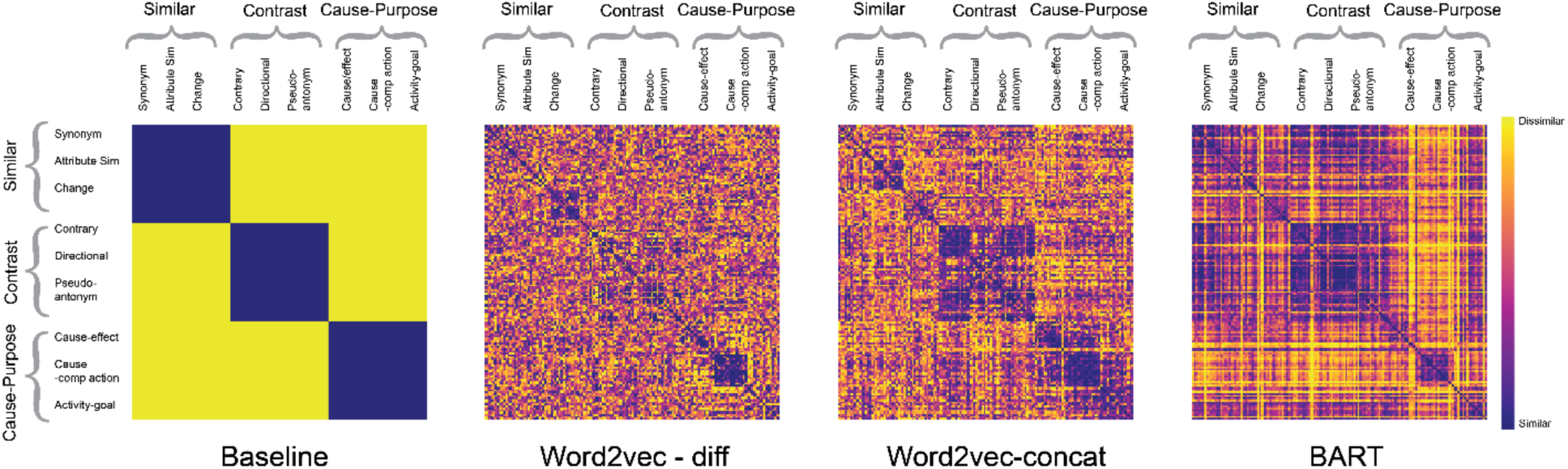
Theoretical Representational Dissimilarity Matrices (RDMs). The RDMs derived from the three computational models are of size 144 × 144 (i.e., based on individual word pairs). Theoretical RDMs capturing the cosine distance between the vector representation for each word pair were correlated with empirical RDMs derived from brain activity patterns.

**Figure 6.**
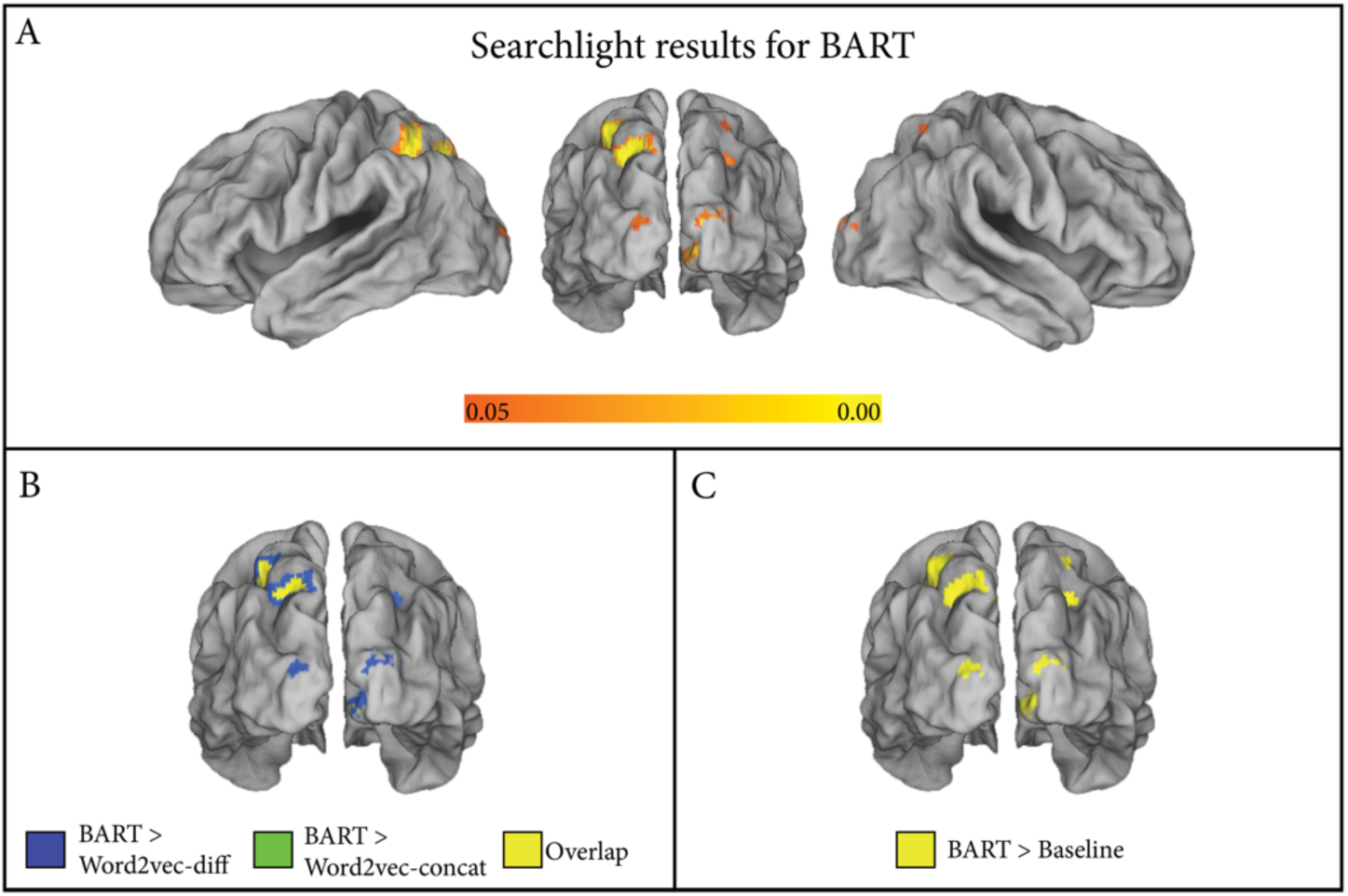
Searchlight results for RSA analyses testing alternative models as predictors of neural similarity during *A:B* phase for 144 word pairs instantiating abstract semantic relations. A: Lateral and posterior views of areas in which the BART model based on distributed relation representations was significantly correlated with neural RDM. None of the other three models yielded areas with significant correlations. B: Posterior view of areas in which correlation of BART with neural RDM was significantly greater than correlation for each of the alternative computational models. C: Posterior view of areas in which correlation of BART with neural RDM was significantly greater than that for the baseline model, which assumes discrete codes for relations. Colored regions represent searchlight sphere centers that were significant as assessed by FSL randomise with TFCE cluster correction (*28, 35*) for multiple comparisons (corrected *p* < 0.05).

### Comparing Computational Accounts of Second-order Relational Processing (C:D Phase)

With respect to the phase in which the *A:B* and *C:D* relations are compared to verify the validity of the analogy (i.e., *C:D* phase), BART and the two Word2vec models make the general prediction that the difficulty of identifying a valid analogy is proportional to the (word or relation-based) similarity of the *A:B* and *C:D* word pairs, with greater dissimilarity making the analogy harder to verify. Valid analogies by definition have the same relation for *A:B* and *C:D*; hence the design-matrix model based on relation-types predicts no differences. In order to derive a measure of *relational dissimilarity* from each of the three models, for every valid *A:B::C:D* analogy (144 problems in total) we calculated the cosine distance between the representations of *A:B* and *C:D* specified by each model, with higher cosine distance implying greater dissimilarity between the two pairs of words. For each individual participant, the theoretical relational dissimilarity scores derived from each model were then correlated (using Spearman’s rho) with observed mean activity during the *C:D* phase of each valid analogy for each region of interest (ROI) (see *SI Appendix*, Univariate Relational Dissimilarity Analysis). ROIs were taken from previous work showing that regions within a left frontoparietal network play prominent roles in relational reasoning (*31*) (see *Materials and Methods*).

As shown in Figure 7, BART’s predictions of theoretical relational dissimilarity were significantly correlated with neural activity in frontal ROIs, with all *p* values FDR-corrected (BA9: mean *r* = 0.06, *p* = 0.022; BA44: *r* = 0.105, *p* = 0.003; BA45: *r* = 0.135, *p* < 0.001; BA10m: *r* = 0.052, *p* = 0.022; BA10l: *r* = 0.069, *p* = 0.006). In addition, the predictions of the BART model, but neither of the alternative ones, were significantly correlated with activity in subregions of parietal cortex (AG: *r* = 0.085, *p* < 0.001; pSMG: *r* = 0.055, *p* = 0.014; and trending in the IPS, *r* = 0.0521, *p* = 0.081).

**Figure 7.**
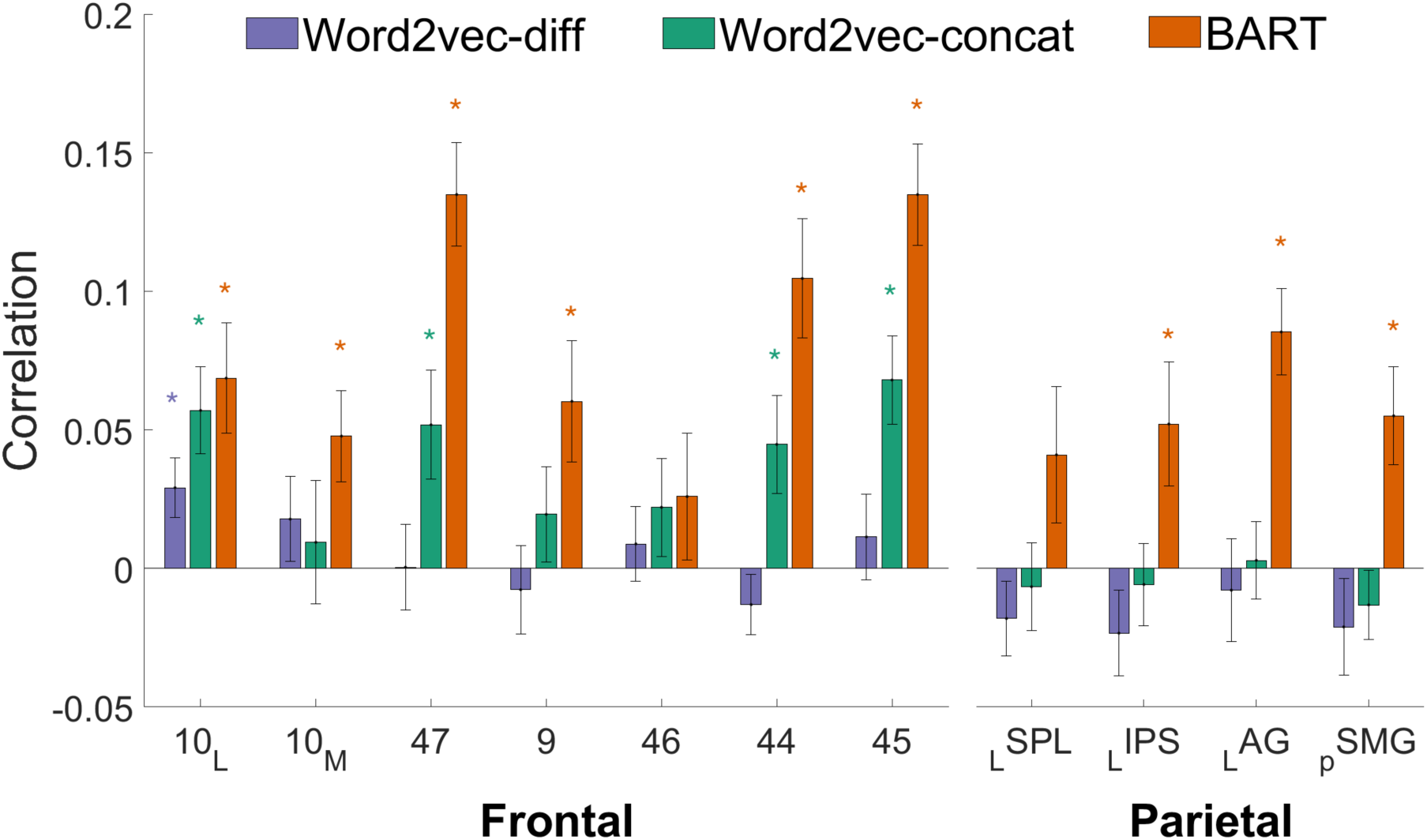
Relation comparison in the brain. Correlation between model-derived relational dissimilarity and mean BOLD signal in each frontal and parietal ROI during the *C:D* phase. Error bars indicate ± 1 standard error of the mean; * indicates model correlation significantly different from 0 (*p* < .05 after FDR correction).

Word2vec-concat’s predictions of relational dissimilarity were correlated with neural activity in some of the same frontal ROIs as BART (BA44: *r* = 0.045, *p* = 0.03; BA45: *r* = 0.068, *p* = 0.003; BA47: *r* = 0.052, *p* = 0.026; BA10l: *r* = 0.057, *p* = 0.009), but not in parietal ROIs. Follow-up stepwise regression analyses (see *SI Appendix*, Supplemental Data Analyses) revealed that in each of the prefrontal ROIs where BART and also Word2vec-concat yielded significant correlations, the BART model explained unique variance in brain activity after accounting for the influence of word meanings as measured by the predictions of the alternative model. After adjusting for Word2vec-concat predictions, semi-partial correlations between BART’s predictions and neural activity were significant in four prefrontal ROIs (BA44: *r* = 0.109, *p* = 0.015, BA45: *r* = 0.136, *p* = 0.005, BA 10l: *r* = 0.07, *p* = 0.01, BA 47: *r* = 0.139, *p* = 0.006). Neural activity in BA10l also showed a correlation with Word2vec-diff predictions. However, the BART model still explained unique variability in brain activity in BA 10l as revealed by a significant semi-partial correlation between BART’s prediction after adjusting for Word2vec-diff prediction (*r* = 0.029, *p* = 0.020). Stepwise regressions with a reversed order revealed that neither Word2vec-concat nor Word2vec-diff predicted variance beyond that attributable to BART.

The coarse semantic coding assumed by the relations-type model is completely unable to explain the fine-grained differences in neural responses observed among the pool of valid analogies during the verification phase of analogical reasoning. In general, the process of second-order relation comparison appears to be dominated by relational dissimilarity as measured by the BART model, which creates distributed representations of relations.

## Discussion

The present study combined computational modeling and neuroimaging to investigate the representation and comparison of abstract semantic relations in the brain. We used sequential presentation of verbal analogies with clear temporal phases (*23*) to decouple (1) the neural activity associated with encoding the representation of the individual words in a pair and the relation between them from (2) that associated with the comparison of two relations (while also separating these high-level reasoning processes from planning for a motor response). By testing alternative computational models of relational similarity, we were able to distinguish between rival accounts of how semantic relations are coded and compared in the brain. The BART model, which postulates that semantic relations between words are coded as a distributed representation based on a taxonomy of abstract relations (*16*), was able to predict patterns of neural activity during analogical reasoning that could not be explained by alternative models. During the phase in which a single relation is being encoded (*A:B* phase), the BART model was the most effective predictor of patterns of neural activity in the superior parietal cortex, a region previously associated with relation representation (*36, 37*)). During the phase in which relations are compared to verify whether the analogy is valid (*C:D* phase), BART was uniquely predictive of neural activity in ROIs spanning the frontal and parietal cortices.

The present findings support three major conclusions. First, semantic relations have distributed representations, primarily coded in subregions of the parietal cortex. Second, the core of these distributed representations is based on a relatively small number of abstract relation types specified in a theory-based taxonomy (*16*); thus we are able to draw inferences not only about the form of relation representations, but also about their specific content. Third, the process of relation representation (*A:B* phase) can be decoupled from the process of making second-order comparisons to solve analogy problems (*C:D* phase).

The fact that similarity measures derived from the BART model yielded stronger and more reliable predictions of relational processing—both of individual relations, and of comparisons between relations—than did the Word2vec-diff model is consistent with computational evidence favoring the former model as an account of human relational judgments (*21*). The relative success of the BART model in predicting patterns of neural activity is directly relevant to a debate as to whether or not individual semantic relations have explicit representations (for discussion see Ref. (*15*)). Whereas Word2vec-diff provides only a generic and implicit representation of relational similarity (i.e., the difference vector between semantic vectors for two words), BART learns representations of individual semantic relations, which then collectively provide an explicit distributed representation of the relations(s) linking any word pair. The neural evidence favoring the BART model of relation similarity thus supports the hypothesis that the brain encodes semantic relations between words as distributed representations across abstract semantic relations, such as *synonym, antonym*, and *cause-effect*. By coupling computational modeling with analyses of similarity in neural activity, it proved possible to resolve theoretical issues that have been debated for the past half century.

The present study focused on abstract semantic relations. These are particularly important because a pool of abstract relations provides basic elements that can be used to represent more specific relations. However, further research will be required to determine the extent to which the neural basis for relational reasoning may differ for more concrete semantic and visuospatial relations (e.g., inferring that grasping a hammer enables it to be lifted). More generally, future studies may benefit from applying the overall strategy of model-guided similarity analyses. This approach has the potential to be used to analyze patterns of neural activity underlying semantic representations of information units more complex than individual words. Careful task design (e.g., presenting a problem in sequential phases) can be used to separate key component processes. Alternative computational models can then be used to generate item-level predictions of neural similarity, which can be tested by methods such as Representational Similarity Analysis. This research strategy shows promise in decoupling component processes and in identifying specific representations underlying high-level reasoning. Future work should aim to develop and test well-specified computational models of how propositions and larger knowledge units are represented in the brain and used to reason.

## Materials and Methods

### Participants

Sixteen participants (8 female) were recruited at the University of California, Los Angeles (UCLA) through a flyer distributed in the Psychology department. Participants signed informed consent prior to the experimental session, and were paid $50 for their participation in the 1-hour study, in compliance with the procedures accepted by the local institutional review board (IRB). The study was approved, including informed consent procedures, by the UCLA Office of the Human Research Protection Program.

### Stimuli

The stimuli were a set of analogy problems constructed from word pairs taken from a normed set of examples of abstract relations (*22*). These norms were in turn based on a linguistic taxonomy of semantic relations (*16*). The full norms include examples of word pairs instantiating ten general types of relations, each including five to ten more specific relations, for a total of 79 distinct relations. For the present study, we focused on three relation types with three specific relations drawn from each, for a total of nine relations: *similar* (*synonym, attribute similarity, change*); *contrast* (*contrary, directional, pseudoantonym*); *cause-purpose* (*cause:effect, cause:compensatory action, activity:goal*). For each relation, we selected 16 word pairs from among the most highly rated (i.e., most prototypical) examples. In making this selection we avoided duplicate pairs that were simple reversals (e.g., *happy-sad* and *sad-happy*), choosing in such cases the pair with the higher typicality rating. Pairs that included conspicuously long or low-frequency words were also excluded. Because for some subcategories it proved difficult to identify 16 pairs that passed our selection criteria, we also included some pairs that (*22*) had used as “seed” examples to elicit word pairs from humans. These were considered excellent examples (most taken from (*16*). The full list of word pairs is provided in the *SI Appendix*, Supplemental Materials and Methods, Table 4.

Using the 144 (16 examples × 9 specific relations) distinct word pairs selected as described above, we formed pairs of pairs to create verbal analogy problems in the form *A:B* :: *C:D* (valid) or else *A:B* :: *C’:D’* (invalid), where all pairs were drawn from the pool of 144. For the invalid pairs, the *C’:D’* pair was drawn from a different relation type than was *A:B*. We avoided creating invalid items using different specific relations within the same general relation type (e.g., specific relations *contrary* and *pseudoantonym*, both subtypes of *contrast*) because pilot work suggested that such “near-miss” problems would lead to excessive errors. At the same time, *C’:D’* pairs always instantiated a natural semantic relation (rather than being semantically anomalous), forcing participants to consider the paired relations carefully in judging validity of the analogies.

Counterbalancing was used to create four complete sets of analogy problems. To form an individual set, for each of the nine specific relations, eight of the 16 pairs were assigned to the *A:B* role and four to the *C:D* role. The remaining four pairs were assigned to the *C’:D’* role associated with *A:B* pairs for four of the six specific relations representing the two remaining general relation types. Assignments to the *C:D* role were random subject to the above restriction. Subject to all of the above restrictions, specific 4-term analogy problems were created by random pairing of word pairs. Each set thus consisted of 72 analogy problems (9 specific relations × 8 problems each). For each specific relation, four problems were valid and four were invalid. Within a set of 72 problems, each of the 144 word pairs occurred twice in the *A:B* role and once in each of the *C:D* and *C’:D’* roles. The same procedure was used to create a total of four sets, each with 72 problems distributed as described above. Across all four sets, each of the 144 word pairs appeared in each role with the same proportions (i.e., twice as often as *A:B* than as *C:D* or C’D’). The four sets, with a total of 288 problems (4 sets × 72 problems each), were treated as four blocks administered to each participant. The procedure for problem generation ensured that any individual analogy problem occurred only once in the set of 288 problems. The order of problems was randomized within each block, and the order of the four blocks was counterbalanced across participants. The overall aim of this procedure for problem creation was to ensure that data analyses could be based on neural patterns associated with each of the 16 word pairs representing each of the nine specific relations (144 pairs in total), in each of the three possible roles (*A:B, C:D, C’:D’*), while avoiding any confounding between specific pairs and roles. Finally, each of these four sets was further split into two sets of 36 for presentation convenience.

### Procedure

The experiment was administered using PsychoPy2 (*38*). On each trial (see Figure 2), participants were first shown the *A:B* word pair for 2s, then the *C:D* pair for 2s (with an average .5s jitter in between). The words “yes” or “no” then appeared on the left and right of the screen, indicating the assignment of two response buttons used to indicate whether or not the two pairs represented the same relation. Critically, the assignment of “yes” and “no” buttons was randomly varied, ensuring that participants could not begin planning a motor response during the earlier phases of the trial.

### fMRI Data Acquisition

Data were acquired on a 3 Tesla Siemens Prisma Magnetic Resonance Imaging (MRI) scanner at the Staglin IMHRO Center for Cognitive Neuroscience at UCLA. Structural data were acquired using a T1-weighted sequence (MPRAGE, TR = 1,900 ms, TE = 2.26 ms, voxel size 1 mm^3^ isovoxel). Blood oxygenation level dependent (BOLD) data were acquired with a T2*-weighted Gradient Recall Echo sequence (TR = 1,000 ms, TE = 37 ms, 60 interleaved slices (2mm gap), voxel size 2×2×2 mm, 6× multiband acceleration).

### fMRI Preprocessing

Data preprocessing was carried out using FSL (*39*). Prior to univariate analyses, data underwent preprocessing steps including motion correction, slice-timing correction (using Fourier-space time-series phase-shifting), spatial smoothing using a Gaussian kernel of 5 mm full-width half-max, and highpass temporal filtering (Gaussian-weighted least-squares straight line fitting, with s=50.0s). Data from each individual run were analyzed employing a univariate general linear model approach (*40*) inclusive of a pre-whitening correction for autocorrelation.

Spatial smoothing was omitted from the above preprocessing steps for classification and Representational Similarity Analysis in order to preserve spatial heterogeneities. Beta-series (*41*) parameter estimates were derived using the Least Squares-Separate, LS-S approach (*42*). The LS-S algorithm iteratively estimates parameters for each trial using a general linear model including a regressor for that trial as well as another regressor for all other trials.

### ROI Selection

For the univariate analysis of neural activity during the *C:D* phase, we selected a number of ROIs associated with relational reasoning within the left lateral frontoparietal network based on meta-analyses of studies of relational reasoning (*30, 31, 37, 43*). These ROIs were defined using the Juelich, Sallet, Neubert, and Harvard Oxford atlases in FSL (*44*). The variety of atlases was used so ROIs would be roughly the same size, and so that ROIs would cover previously-reported coordinates based on the relevant meta-analyses. ROIs included BA10, which was separated into a medial BA10 (BA10m) defined by the Sallet dorsal frontal connectivity parcellation, and a lateral BA10 (BA10l) defined by the Neubert ventral frontal connectivity parcellation. These two ROIs, together with BA47 (see below), were selected to fully cover the area of previously-reported activations in rlPFC. We also selected areas from the ventrolateral and dorsolateral PFC, including BA9 (defined using the Sallet frontal connectivity parcellation), BAs 44 and 45 (defined by the Harvard Oxford atlas), and BA 47 (selected using the Neubert frontal connectivity parcellation). In the parietal cortex, we used the Juelich histological atlas. We created the IPS ROI by taking the union of all IPS subdivisions (*37*). We also selected the superior parietal lobe (SPL 7A), and two subdivisions of the inferior parietal lobe (PFm and PGa) corresponding to the angular gyrus (AG) and posterior supramarginal gyrus (pSMG).

## Data Availability

Raw and preprocessed NIFTI files, as well as experiment timing files will be uploaded to a repository (openfmri.org), and is available from the first author upon request.

## Code Availability

Code for the BART model can be downloaded from cvl.psych.ucla.edu/BART2code.zip. Code for the experiment and all custom analyses can be found at https://github.com/njchiang/analogy-fmri.

## Acknowledgements

Preparation of this paper was supported by NSF Grant BCS-1827374 to H.L. and K.J.H., a UCLA Academic Senate grant to K.J.H., and a Department of Defense (DoD) grant through the National Defense Science and Engineering Graduate Fellowship Program to J.N.C. A preliminary report of this work was presented at the 2018 meeting of the Society for Neuroscience (San Diego, November).

## Author Contributions

Conceptualization, K.J.H, H.L., and M.M.M.; Methodology, J.N.C., Y.P., H.L., and M.M.M.; Formal Analysis, J.N.C. and H.L.; Investigation, J.N.C. and Y.P.; Writing – Original Draft, J.N.C. and K.J.H.; Writing – Review & Editing, J.N.C., Y.P., H.L., K.J.H., and M.M.M.; Supervision, M.M.M; Funding Acquisition, H.L. and K.J.H.

## Declaration of Interests

The authors declare no competing interests.

## Notes

#### Summary of Updates

Major manuscript revision

