## Supplementary material for "Distributed Code for Semantic Relations Predicts Neural Similarity"

**Supporting material for Chiang et al.**

### I. Supplemental Data Analyses

#### Behavioral Results

Mean proportion correct in solving analogy problems was 0.82 (*SD* = 0.07) across all conditions, an accuracy level well above chance ($p < .001$). A repeated-measures ANOVA was conducted on performance accuracy across the three abstract relation types for *A:B*. Problems using the *contrast* relation yielded the highest accuracy (*M* = 0.87, *SD* = 0.08), followed by *cause-purpose* (*M* = 0.82, *SD* = 0.07) and *similar* (*M* = 0.78, *SD* = 0.09). Bonferroni-corrected *t* tests indicated that *contrast* problems were more accurate than either of the other relation types ($p < .005$).

Reaction time was calculated as the time from the appearance of the response cue (i.e., "yes" and "no" indicators after the *C:D* phase) to the button press. Only reaction times for accurate trials were analyzed. Mean reaction time was 913 ms (*SD* = 255 ms) across all relation types. No reliable reaction time differences were found between the three types.

#### Multivariate Analyses: Areas that Distinguish Among Abstract Semantic Relations

To characterize the representations of abstract semantic relations in the brain, we conducted a multivariate classification analysis using a searchlight method (Kriegeskorte, Goebel, & Bandettini, 2006). Classifiers were trained to distinguish between the three general relation types (*similar*, *contrast*, *cause-purpose*), and were evaluated using a leave-one-run-out cross-validation approach (see Etzel & Braver, 2013). For each participant, two such classifications were run: one on the *A:B* phase and one on the *C:D* phase (including both valid and invalid trials). We used a 5mm radius sphere and a linear SVM (Abraham et al., 2014; Pedregosa et al., 2011). Statistical significance was assessed using FSL randomise with TFCE cluster correction (Smith & Nichols, 2009; Winkler, Ridgway, Webster, Smith, & Nichols, 2014).

This analysis revealed brain areas capable of distinguishing different types of abstract relations on the basis of activation patterns during the *A:B* and *C:D* phases. As shown in Figure 3 in main paper, distributed areas of the brain are involved in decoding semantic relation types (*similar*, *contrast*, and *cause-purpose*). Areas in color achieved above-chance classification performance ($p < 0.01$), as assessed by a Wilcoxon signed-rank test with TFCE cluster correction (Smith & Nichols, 2009). During the *A:B* phase, the active regions for relation classification include frontal and temporal cortices (most pronounced in the left hemisphere), and bilateral parietal cortices. During the *C:D* phase, the three relation types can also be distinguished in many of the same regions, and also in additional regions in the right hemisphere (particularly across frontal and temporal cortices). Overall, the overlap in regions capable of distinguishing the three semantic relations across both *A:B* and *C:D* phases (areas in yellow in Figure 3) includes areas previously suggested to be part of the semantic representation system for individual words (Binder, Desai, Graves, & Conant, 2009; Carota, Kriegeskorte, Nili, & Pulvermüller, 2017; de Heer, Huth, Griffiths, Gallant, & Theunissen, 2017; Huth, de Heer, Griffiths, Theunissen, & Gallant, 2016). The analysis also highlights the important role of parietal regions associated more specifically with relational reasoning (Wendelken, 2015).

**Univariate Analyses: Localization of Relation Representation and Comparison**

The general relation type was coded separately for the *A:B* and *C:D* phases of each trial (including both valid and invalid trials). A univariate analysis using the GLM approach was performed to identify regions engaged in representing semantic relations. The response phase of each trial was included as a condition of non-interest, as well as motion parameters. The GLM analysis was carried out using FSL FEAT (Jenkinson, Beckmann, Behrens, Woolrich, & Smith, 2011; S. M. Smith et al., 2004). Data from individual runs were aggregated employing a mixed effects model (i.e., employing both the within- and between-subject variance), and using automatic outlier detection. Statistical significance for univariate analyses were assessed using FSL randomize with TFCE cluster correction (Smith & Nichols, 2009; Winkler et al., 2014). The following contrasts are displayed in Figure S1: (left) *A:B* – rest, *C:D* – rest; (right) *C:D* – *A:B*.

As shown in Figure S1 (left), in the *A:B* stage, related word pairs elicited mostly left-lateralized frontal and temporal activity, bilateral parietal activity, and activity in the occipital lobe (see Table S1 for detailed list). The *C:D* stage, compared to simple fixation, recruited many of the same regions as did the *A:B* stage (likely involved in processing each word of the *C:D* pair and encoding their semantic relation), as well as unique activations likely involved in second-order relation assessment for relation comparison. Specifically, the *A:B* and *C:D* stimuli shared activations in the inferior lateral occipital cortex (BA19), fusiform gyrus (BA37), and left frontal regions spanning the rostrolateral prefrontal cortex (BA10 and BA47). In addition, processing *C:D* word pairs uniquely led to a greater BOLD response in left inferior frontal gyrus (*pars triangularis, pars opercularis;* BA44, 45) as well as bilateral superior parietal cortex (in BA7; for a full list see Table S2).

As shown in Figure S1 (right), the univariate comparison of *C:D* and *A:B* phases revealed a fronto-parietal network, mainly left lateralized, presumably involved in the process of second-order relation comparison, which recruits additional regions beyond those related to processing individual words and their semantic relation(s). Specifically, the contrast uncovered significant clusters in the left RLPFC (BAs 10, 47), replicating prior results implicating this region in complex relational comparisons (Bunge, Helskog, & Wendelken, 2009; Christoff et al., 2001), as well as in the left inferior frontal gyrus (BAs 44 and 45), bilateral posterior parietal (BA7) and occipital cortices (BA19) (for a full list see Table S3).

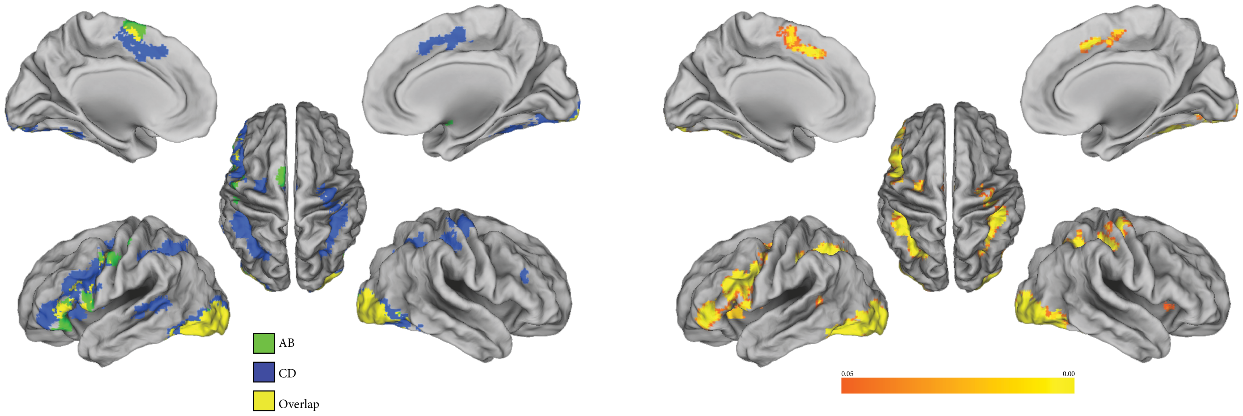


*Figure S1*. Univariate analysis results. Left: Main effects of *A:B* and *C:D* phases of trials. Clusters were obtained by contrasting each phase (i.e., *A:B*, *C:D*) to simple fixation. Right: *C:D* – *A:B* univariate contrast. Regions in which activity while reading the *C:D* word pair was greater than when reading the *A:B* word pair. Depicted group-level activations were obtained with a non-parametric permutation approach (FSL randomize), significance was set at $p$=0.05 FWER, cluster-corrected with threshold-free cluster enhancement (TFCE; Smith & Nichols, 2009).

Table S1: Local maxima for the univariate contrast of *A:B* phase versus rest.

| Lobe | Coord (MNI) x | y | z | Z | Hem | Anatomical Label |
|---|---|---|---|---|---|---|
| Frontal | | | | | | |
| | -54 | 14 | 2 | 4.67 | L | Inferior Frontal Gyrus (pars opercularis) |
| | -50 | 10 | 4 | 4.66 | L | Inferior Frontal Gyrus (pars opercularis) |
| | -4 | 2 | 64 | 5.2 | L | Juxtapositional Lobule Cortex (SMA) |
| | 0 | 8 | 70 | 4.55 | | Juxtapositional Lobule Cortex (SMA) |
| | -44 | 28 | -12 | 4.86 | L | Orbitofrontal Cortex |
| | -50 | -6 | 42 | 4.9 | L | Precentral Gyrus |
| | -52 | -4 | 48 | 4.68 | L | Precentral Gyrus |
| | -6 | -4 | 72 | 3.38 | L | Superior Frontal Gyrus |
| | 20 | -2 | 14 | 3.69 | R | WM |
| Occipital | | | | | | |
| | -34 | -88 | -4 | 5.98 | L | Lateral Occipital Cortex (inferior division) |
| | -36 | -86 | -8 | 5.93 | L | Lateral Occipital Cortex (inferior division) |
| | -40 | -86 | -6 | 5.81 | L | Lateral Occipital Cortex (inferior division) |
| | -46 | -70 | -18 | 5.43 | L | Lateral Occipital Cortex (inferior division) |
| | 48 | -64 | -22 | 5.25 | R | Occipital Fusiform Gyrus |
| | -34 | -92 | -2 | 5.7 | L | Occipital Pole |
| | -30 | -96 | -2 | 5.48 | L | Occipital Pole |
| | 22 | -92 | -12 | 6.55 | R | Occipital Pole |
| | 32 | -90 | -12 | 6.47 | R | Occipital Pole |
| | 28 | -90 | 0 | 6.08 | R | Occipital Pole |
| | 22 | -96 | -6 | 5.5 | R | Occipital Pole |
| | 24 | -94 | 0 | 5.33 | R | Occipital Pole |
| Subcortical | | | | | | |
| | 26 | 2 | -14 | 3.82 | R | Amygdala |
| | 16 | 10 | 4 | 4.12 | R | Caudate |
| | 20 | 10 | 14 | 3.83 | R | Caudate |
| | -20 | 10 | 10 | 5.63 | L | Putamen |
| | 24 | 4 | 2 | 4.26 | R | Putamen |
| | 28 | 12 | -2 | 3.71 | R | Putamen |

Table S2: Local maxima for the univariate contrast of *C:D* phase versus rest.

| Lobe | Coord (MNI) x | y | z | Z | Hem | Anatomical Label |
|---|---|---|---|---|---|---|
| Frontal | | | | | | |
| | -48 | 40 | 4 | 6.29 | L | Frontal Pole |
| | 30 | 40 | 14 | 3.33 | R | Frontal Pole |
| | -44 | 16 | 28 | 6.43 | L | Inferior Frontal Gyrus (pars opercularis) |
| | -44 | 18 | 24 | 5.75 | L | Inferior Frontal Gyrus (pars opercularis) |
| | 30 | 30 | 16 | 3.83 | R | Inferior Frontal Gyrus (pars triangularis) |
| | 28 | 18 | 12 | 3.4 | R | Insula |
| | 38 | 28 | 24 | 4.1 | R | Middle Frontal Gyrus |
| | 22 | 18 | 20 | 3.71 | R | None |
| | 28 | 36 | 14 | 3.58 | R | None |
| | -42 | 6 | 30 | 6.16 | L | Precentral Gyrus |
| | 28 | -14 | 52 | 5.88 | R | Precentral Gyrus |
| | 30 | -12 | 56 | 5.62 | R | Precentral Gyrus |
| Occipital | | | | | | |
| | -36 | -84 | -8 | 7.45 | L | Lateral Occipital Cortex (inferior division) |
| | -30 | -90 | -4 | 7.09 | L | Lateral Occipital Cortex (inferior division) |
| | 28 | -86 | -8 | 7.66 | R | Occipital Fusiform Gyrus |
| | 22 | -92 | -12 | 8.36 | R | Occipital Pole |
| | 32 | -90 | -12 | 7.84 | R | Occipital Pole |
| | 22 | -94 | -6 | 7.75 | R | Occipital Pole |
| Parietal | | | | | | |
| | -24 | -62 | 40 | 5.32 | L | Lateral Occipital Cortex (superior division) |
| | -28 | -60 | 40 | 5.15 | L | Lateral Occipital Cortex (superior division) |
| | -26 | -64 | 46 | 5.03 | L | Lateral Occipital Cortex (superior division) |
| | -26 | -64 | 34 | 4.94 | L | Lateral Occipital Cortex (superior division) |
| | -26 | -50 | 38 | 6.62 | L | Superior Parietal Lobule |
| | -40 | -40 | 42 | 5.86 | L | Supramarginal Gyrus (anterior division) |

Table S3: Local maxima for the univariate *C:D* - *A:B* contrast.

| Lobe | Coord (MNI) x | y | z | Z | Hem | Anatomical Label |
|---|---|---|---|---|---|---|
| Frontal | | | | | | |
| | 32 | 28 | 14 | 3.36 | R | Frontal Operculum Cortex |
| | -44 | 40 | 2 | 5.21 | L | Frontal Pole |
| | 32 | 42 | 14 | 3.28 | R | Frontal Pole |
| | -42 | 16 | 26 | 5.53 | L | Inferior Frontal Gyrus (pars opercularis) |
| | -46 | 20 | 22 | 4.9 | L | Inferior Frontal Gyrus (pars opercularis) |
| | 36 | 26 | 18 | 3.42 | R | Inferior Frontal Gyrus (pars triangularis) |
| | -46 | 24 | 24 | 5.16 | L | Middle Frontal Gyrus |
| | 40 | 30 | 24 | 3.58 | R | Middle Frontal Gyrus |
| | 48 | 34 | 26 | 3.45 | R | Middle Frontal Gyrus |
| | 42 | 34 | 20 | 3.4 | R | Middle Frontal Gyrus |
| | -42 | 8 | 28 | 4.98 | L | Precentral Gyrus |
| | -44 | 6 | 32 | 4.86 | L | Precentral Gyrus |
| | 42 | -2 | 14 | 4.3 | R | Central Opercular Cortex |
| | 42 | 2 | 14 | 4.08 | R | Central Opercular Cortex |
| | 48 | -4 | 12 | 3.51 | R | Central Opercular Cortex |
| | 40 | -2 | 20 | 3.27 | R | Central Opercular Cortex |
| | 38 | -20 | 22 | 3.52 | R | Parietal Operculum Cortex |
| | 38 | -24 | 20 | 3.32 | R | Parietal Operculum Cortex |
| Occipital | | | | | | |
| | 38 | -84 | -2 | 5.75 | R | Lateral Occipital Cortex (inferior division) |
| | -48 | -68 | -6 | 5.72 | L | Lateral Occipital Cortex (inferior division) |
| | 40 | -80 | 2 | 5.63 | R | Lateral Occipital Cortex (inferior division) |
| | 48 | -76 | 4 | 5.63 | R | Lateral Occipital Cortex (inferior division) |
| | 20 | -96 | 4 | 8.45 | R | Occipital Pole |
| | 16 | -92 | -14 | 7.41 | R | Occipital Pole |
| | 36 | -62 | 54 | 6.06 | R | Lateral Occipital Cortex (superior division) |
| | 24 | -58 | 40 | 5.56 | R | Lateral Occipital Cortex (superior division) |
| | -30 | -72 | 48 | 5.61 | L | Lateral Occipital Cortex (superior division) |
| | -18 | -70 | 44 | 4.46 | L | Lateral Occipital Cortex (superior division) |
| | -22 | -66 | 46 | 4.46 | L | Lateral Occipital Cortex (superior division) |
| Parietal | | | | | | |
| | 2 | 8 | 48 | 5.29 | R | Paracingulate Gyrus |
| | 40 | -28 | 56 | 5.59 | R | Postcentral Gyrus |

| | | | | | | |
|---|---|---|---|---|---|---|
| | 32 | -28 | 50 | 5.34 | R | Postcentral Gyrus |
| | 30 | -12 | 52 | 5.17 | R | Precentral Gyrus |
| | -26 | -50 | 40 | 4.94 | L | Superior Parietal Lobule |
| | -32 | -50 | 44 | 4.74 | L | Superior Parietal Lobule |
| | -40 | -40 | 42 | 5.7 | L | Supramarginal Gyrus (anterior division) |
| Temporal | | | | | | |
| | -62 | -42 | 4 | 5.67 | L | Middle Temporal Gyrus (posterior division) |
| | -52 | -48 | 4 | 4.48 | L | Middle Temporal Gyrus (temporooccipital part) |
| | -64 | -36 | 6 | 5.4 | L | Superior Temporal Gyrus (posterior division) |
| | -48 | -32 | -2 | 4.61 | L | Superior Temporal Gyrus (posterior division) |
| | -50 | -40 | 8 | 4.56 | L | Superior Temporal Gyrus (posterior division) |
| | -52 | -46 | 10 | 4.68 | L | Supramarginal Gyrus (posterior division) |

### Distinguishing Valid Versus Invalid Trials

An ROI classification analysis was run to determine which regions encoded information meaningful to solving the analogy (Figure S2). Using a leave-one-run-out cross-validation procedure (cf. Etzel & Braver, 2013), a classifier was trained to discriminate between valid vs invalid trials based on multivariate ROI activity. Within frontal and parietal ROIs, classifiers were able to distinguish whether subjects were looking at a valid (*C:D*) or invalid (*C':D'*) word pair, whereas within control regions (occipital, CSF) classifers were unable to do so.

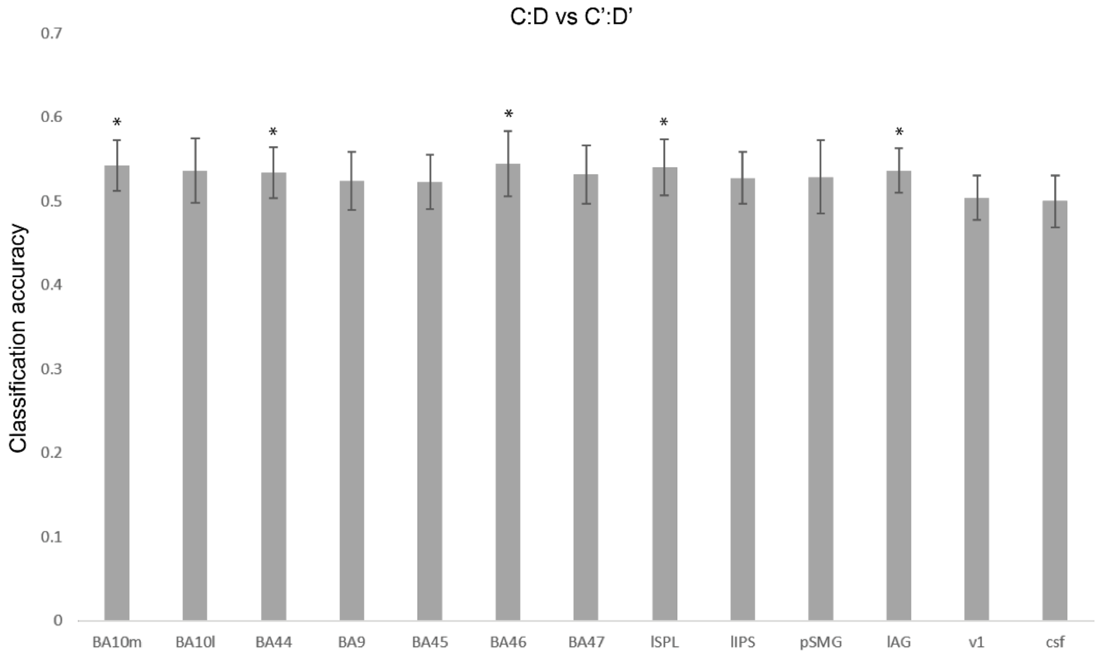


*Figure S2.* ROI classification analysis for distinguishing valid (*C:D*) from invalid (*C':D'*) analogy problems. Error bars represent +/- 1 SEM.

**Representational Similarity Analysis**

Representational Similarity Analysis (RSA; (Kriegeskorte, Mur, & Bandettini, 2008; Kriegeskorte & Kievit, 2013; Nili et al., 2014)was used to characterize the similarities of neural responses across pairs. RSA characterizes the representation in a brain region by a representational dissimilarity matrix (RDM), and compares this empirical matrix with a theoretical model. An RDM is a square symmetric matrix, with each entry referring to the dissimilarity between the activity patterns associated with two trials (e.g., entry (1,2) would represent the dissimilarity between activity patterns on trial 1 and trial 2). Procedurally, each element of the RDM is calculated as 1 minus the Pearson correlation between the beta-series for each pair of trials (Carota et al., 2017; Nili et al., 2014).

Hypothesis models were manually generated to reflect idealized RDMs expected given a theoretical representational space. We generated theoretical RDMs from each of the three computational models. Each model uses a different calculation to yield a feature vector characterizing a word pair; however, the RDM was calculated in the same way for all models, as the cosine distance between word-pair representations.

RDMs and hypothesis models were compared by calculating a "second-order similarity" (Nili et al., 2014), defined as the Spearman correlation coefficient between the two matrices. All analyses were carried out using Python, making extensive use of the machine learning packages Scikit-learn (Pedregosa et al., 2011) and NiLearn (Abraham et al., 2014).

**Univariate Relational Dissimilarity Analysis**

All the models of analogical comparison considered in the present paper make the general prediction that the difficulty of deciding the validity of an analogy will be related to the relation-based similarity of the *A:B* and *C:D* word pairs, with greater similarity making the decision easier. In this analysis, only trials consisting of valid analogies (i.e., *A:B* :: *C:D*) were included so that the relation representations during the *C:D* phase would not be confounded by additional cognitive operations associated with processing a relation inconsistent with *A:B*. To derive a specific prediction from each of the three candidate models in the paper, for every valid analogy of word pairs, *A:B::C:D*, a *relational dissimilarity* measure was calculated by taking the cosine distance between the representations of *A:B* and of *C:D* specified by the model (i.e., higher cosine distance implies greater dissimilarity between the two pairs). These model-derived relational dissimilarity scores for each trial were then correlated (using Spearman's rho) with mean ROI activity to identify brain regions that track relational dissimilarity according to the predictions from each of the alternative models. The resulting *p* values were adjusted for multiple comparisons by controlling the false discovery rate (FDR) at q = 0.05 and are reported in the main text.

***C:D* Phase: Follow-up Regression Analysis**

Within frontal and parietal regions that were significantly correlated with both BART and Word2vec-concat models in the *C:D* phase (Figure 6), the general trend was that BART-derived predictions showed greater correlation with ROI-derived relational similarity than with Word2vec-concat, the model based on word similarity. In these ROIs (BA44, BA45, BA47, BA10l), a semipartial correlation analysis was performed to determine whether these two models captured the same or different information. Word2vec-concat relational dissimilarity scores were first regressed out of the ROI-based similarity scores, and subsequently the resulting residuals were correlated with the relational dissimilarity predictions from BART. (See main text for results.) The reverse analysis was also performed, in which BART relational dissimilarity predictions were regressed out of ROI-based dissimilarity scores and then correlated with the

Word2vec-concat predictions. No regions showed a significant impact of Word2vec-concat after controlling for the variance predicted by BART. That is, Word2vec-concat did not appear to capture any information about relational dissimilarity beyond that accounted for by the BART model. The associated $p$ values were corrected for the false discovery rate at $q = 0.05$ and the adjusted values are reported in the main text of the paper.

### II. Supplemental Materials and Methods

#### Counterbalancing Details

Within a set of 72 problems (see *Materials and Methods*, Stimuli), each of the 144 word pairs occurred twice in the *A:B* role and once in each of the *C:D* and *C':D'* roles. Across the four sets of problems, each of the 144 word pairs appeared in each role with the same proportions (i.e., twice as often as *A:B* than as *C:D* or C'D'). The four sets, with a total of 288 problems (4 sets x 72 problems each), were treated as four blocks administered to each participant. The procedure for problem generation ensured that any individual analogy problem occurred only once in the set of 288 problems. The order of problems was randomized within each block, and the order of the four blocks was counterbalanced across participants. The overall aim of this procedure for problem creation was to ensure that data analyses could be based on neural patterns associated with each of the 16 word pairs representing each of the nine specific relations (144 pairs in total), in each of the three possible roles (*A:B*, *C:D*, *C':D'*), while avoiding any confounding between specific pairs and roles. Finally, each of these four sets was further split into two sets of 36 for presentation convenience.

Table S4: Word pairs used to generate analogy problems, organized by major type and subtype.

| | *Similar* | |
|---|---|---|
| *Synonym* | *Attribute Similarity* | *Change* |
| big:large | book:magazine | acceleration:speed |
| boat:ship | chair:sofa | darken:color |
| car:auto | fence:hedge | death:population |
| careful:cautious | hill:mountain | dim:light |
| couch:sofa | house:tent | discount:price |
| cute:adorable | ladder:stairs | food:water |
| house:home | paper:parchment | force:pressure |
| kid:child | pencil:pen | heat:temperature |
| make:manufacture | picture:drawing | ination:price |
| option:choice | pillow:cushion | lower:volume |
| pants:trousers | rake:fork | raise:salary |
| pretty:beautiful | shovel:spoon | rise:tide |
| raise:elevate | stairs:ladder | shorten:distance |
| run:sprint | sword:knife | soften:voice |
| spin:twirl | table:desk | speed:movement |
| teach:instruct | wagon:trailer | terror:fear |

| | *Contrast* | |
|---|---|---|
| *Contrary* | *Directional* | *Pseudoantonym* |
| accept:reject | ahead:behind | bright:dull |
| big:small | below:above | day:evening |
| black:white | climb:descend | enthusiastic:lazy |
| bright:dark | east:west | fun:boring |
| dark:light | forward:backward | funny:serious |
| di_cult:easy | front:back | good:wrong |
| dirty:clean | high:low | high:down |
| fast:slow | in:out | just:unfair |
| fat:thin | interior:exterior | loud:discreet |
| good:bad | left:right | low:up |
| hot:cold | north:south | majority:small |
| old:young | rise:sink | obey:protest |
| pretty:ugly | start:finish | powerful:meek |
| rich:poor | top:bottom | right:bad |
| tall:short | under:over | smiling:sad |
| warm:cool | up:down | witty:dumb |

*Cause-purpose*

| *Cause:effect* | *Cause:compensatory action* | *Activity:goal* |
|---|---|---|
| accident:damage | anger:yell | advertise:promote |
| bath:cleanliness | coldness:shiver | bathe:clean |
| disease:sickness | danger:ee | breathe:live |
| exercise:fitness | dirtiness:bathe | burnish:shine |
| explosion:damage | dirty:bathe | cook:eat |
| _re:burns | fright:scream | drink:hydrate |
| germs:sickness | happiness:smile | exercise:healthy |
| heater:warmth | heat:sweat | ee:escape |
| illness:discomfort | hunger:eat | ignite:burn |
| injury:pain | loneliness:socialize | read:learn |
| joke:laughter | nervousness:sweat | sleep:rest |
| loss:grief | sadness:cry | speak:express |
| repetition:boredom | sickness:medicate | study:learn |
| stimulus:response | thirst:drink | trim:shorten |
| tragedy:tears | thirsty:drink | wash:clean |
| workout:sweat | tiredness:rest | work:earn |

### III. Supplemental Details for Computational Models

All quantitative models used to create theoretical RDMs for the RSA analysis were based directly (Word2vec-concat, Word2vec-diff) or indirectly (BART) on the outputs of a machine learning model, Word2vec (Mikolov, Sutskever, Chen, Corrado, & Dean, 2013). This model takes a large text corpus (Google News) as input, examines distributional statistics relating each word to neighboring words in sentences (local context), and outputs a modular vector representation for each individual word, termed a *word embedding*. Word2vec vectors of length 300 were obtained for all words used in the present study. Word2vec-concat (the concatenation of the vectors for the two words in a pair) and Word2vec-diff (the difference vector derived from the two individual vectors) were calculated and used to create theoretical RDMs.

The BART model (*Bayesian Analogy with Relational Transformations*; Lu, Wu, & Holyoak, 2019) was applied to learn specific relations between word pairs. BART takes as inputs pairs of positive and negative examples of a given relation, where each pair is represented by the concatenation of the Word2vec vector for each word. For example, a vector formed by concatenating the individual vectors for *love* and *hate* would constitute a positive example of the *antonymy* relation, but a negative example of the *category membership* relation. The model used supervised learning with 20 positive examples and a fixed set of 64-74 negative instances (the top example for each relation from each general category other than that of the target relation) to form weight distributions representing each of the 79 relations in a set of norms (Jurgens, Turney, Mohammad, & Holyoak, 2012). For each word pair used in the study, these learned weights were used to calculate the posterior probability that the pair instantiated each of the 79 learned relations. The vector of length 79 formed by these posterior probabilities represented the specific relation between the two words in the pair. These vectors were used to create BART's theoretical RDMs.

**References (Specific to SI Appendix)**

Abraham, A., Pedregosa, F., Eickenberg, M., Gervais, P., Mueller, A., Kossaifi, J., … Varoquaux, G. (2014). Machine learning for neuroimaging with scikit-learn. *Front Neuroinform*, *8*, 14. https://doi.org/10.3389/fninf.2014.00014

Carota, F., Kriegeskorte, N., Nili, H., & Pulvermüller, F. (2017). Representational Similarity Mapping of Distributional Semantics in Left Inferior Frontal, Middle Temporal, and Motor Cortex. *Cerebral Cortex*, *27*(1), 294–309. https://doi.org/10.1093/cercor/bhw379

Christoff, K., Prabhakaran, V., Dorfman, J., Zhao, Z., Kroger, J. K., Holyoak, K. J., & Gabrieli, J. D. E. (2001). Rostrolateral Prefrontal Cortex Involvement in Relational Integration during Reasoning. *NeuroImage*, *14*(5), 1136–1149. https://doi.org/10.1006/nimg.2001.0922

de Heer, W. A., Huth, A. G., Griffiths, T. L., Gallant, J. L., & Theunissen, F. E. (2017). The Hierarchical Cortical Organization of Human Speech Processing. *The Journal of Neuroscience*, *37*(27), 6539–6557. https://doi.org/10.1523/JNEUROSCI.3267-16.2017

Etzel, J. A., & Braver, T. S. (2013). MVPA Permutation Schemes: Permutation Testing in the Land of Cross-Validation. *2013 International Workshop on Pattern Recognition in Neuroimaging*, 140–143. https://doi.org/10.1109/PRNI.2013.44

Huth, A. G., de Heer, W. A., Griffiths, T. L., Theunissen, F. E., & Gallant, J. L. (2016). Natural speech reveals the semantic maps that tile human cerebral cortex. *Nature*, *532*(7600), 453–458. https://doi.org/10.1038/nature17637

Pedregosa, F., Varoquaux, G., Gramfort, A., Michel, V., Thirion, B., Grisel, O., … Cournapeau, D. (2011). Scikit-learn: Machine Learning in Python. *Journal of Machine Learning Research*, 6.

Smith, S. M., Jenkinson, M., Woolrich, M. W., Beckmann, C. F., Behrens, E. J., H, J.-B., … Matthews, P. M. (2004). Advances in functional and structural MR image analysis and implementation as FSL. *Neuroimage*, *23*, S208–S219.
